# Marked change of cuticular structural color caused by knocking down of *Distal-less* gene in a leaf beetle

**DOI:** 10.64898/2026.11.23.768737

**Authors:** Masako Katsuki, Takashi Matsuo, Ken Ishikawa, Yasukazu Okada, Manabu Kamimura

## Abstract

Structural color is found on a variety of traits, including body surface and feathers across organisms. Many insect species also exhibit structural coloration in their body cuticle, particularly in Coleoptera and Lepidoptera. Recent studies using SEM and TEM observations have revealed the nanostructures responsible for structural color. However, the developmental and genetic mechanisms underlying the formation of these structures remain largely unknown. Here, we showed that *Distal-less* (*Dll*), a major limb-patterning gene, relates cuticular structural color in the leaf beetle *Sagra femorata.* During post-eclosion development, individuals exhibited a shift in the peak reflectance of structure color from short to long wavelengths. In contrast, *Dll* knocked down individuals retained short wavelength reflectance and did not undergo this shift. From SEM observation, we found interference layers responsible for structural coloration immediately beneath the cuticle surface. Comparison of the reflective layers between control and the *Dll* knocked down individuals revealed differences in layer thickness; electron-lucent layers were thinner and electron-dense layers were thicker in *Dll* knocked down individuals. These results suggest that the primarily limb-patterning gene *Dll* is deployed during cuticle development and is involved in the formation of reflective layers responsible for structural color in *S. femorata*.

## Introduction

The beautiful and brilliant structural colors in organism, such as peacock feathers, shell of mollusks and wings of butterflies, have attracted humans from ancient. In contrast to colors produced by pigments, structural coloration is produced by the physical effect and is found in a variety of traits such as body surface and feathers of organisms spanning from prokaryotes [1–3] to eukaryotes including such as plants [4], insects [5], cephalopods [6] and birds [7].

The physical structures responsible for structural coloration have been revealed through observations using scanning electron microscope (SEM) and transmission electron microscope (TEM) [8–11]. In addition to these observations, mechanisms by which the microstructure acquires the specific structural color have been revealed by physical studies [12–14]. In coleopteran species, structural colors are produced by multilayer interference [15], photonic crystals [16], diffraction gratings [17] and light scattering [18]. Especially, multilayer reflectors in the cuticle have been well reported [5, 19]. Repeated thin layers in the epicuticle or exocuticle reflect specific wavelengths of light, thereby producing structural coloration. The number and thickness of these multilayer reflectors determine the color and brightness of cuticle [15, 20, 21]. Several studies have demonstrated differences in the thickness and number of multilayer reflectors between individuals exhibiting different structural colors [20, 21]. However, the developmental mechanisms and genes responsible for the formation of multilayer reflectors within the cuticle remain largely unknown.

Development of the imago cuticle layers begins from prepupal stage and continues through eclosion [22], involving numerous genes, including those encoding cuticle proteins and enzymes involved in chitin synthesis (reviewed in [23, 24]). Structural color arises from highly ordered multilayer architectures that require precise spatial organization along specific body axes. The formation of such directional layered structures suggests that positional information regulating development along the distal-proximal axis may also contributes to the formation of cuticle reflective layers. Developmental pathways are often co-opted to generate novel morphological traits. Here, we therefore focus on limb patterning genes as candidate involved in forming structural color layers. Limb patterning genes establish positional information along the distal-proximal axis during appendage development. We hypothesize that genes involved in establishing positional information during appendage development may also contribute to the spatial organization of cuticular reflective layers.

The leaf beetle *Sagra femorata* (Drury) (Coleoptera, Chrysomelidae), which has been introduced into Japan, exhibits pink-purple structural coloration. To examine the roles of limb-patterning genes in structural color formation in *S. femorata*, we performed RNA interference (RNAi) targeting three genes, *distal-less* (*Dll*), *dachshund* (*dac*) and *homothorax* (*hth*), which regulate the development of the distal, intermediate and dorsal regions of the appendages, respectively. As the results, we found that *Dll* affects the structural coloration. We then investigated the post-eclosion shift in the wavelength of peak reflectance peak in *Dll* knockdown (KD) and control (*gfp* KD) individuals. Additionally, we compared the surface layers of elytral cuticle between treatments using scanning electron microscopy (SEM) to investigate which region of the cuticle was affected by *Dll* KD. Based on these results, we discuss how silencing of *Dll* gene influences structural color formation in *S. femorata*.

## Materials and Methods

### Insects

*Sagra femorata* larvae develp within the stems of kudzu plant, *Pueraria montana var. lobata* (Leguminosae: Phaseoleae) and overwinter in the gall formed in the stem during the final instar. The larvae used in this study were collected in Mie prefecture in 19th to 21st January 2022, and on 28th February and 1st March 2025. After collecting, all larvae were kept within the galls and stored in a refrigerator until the RNAi experiments.

### Identification of three appendage genes in *Sagra femorata*

Transcript sequences of the *S. femorata* orthologs of *Dll*, *hth* and *dac* were identified from a transcript catalog generated by RNA sequencing (RNA-seq). Total RNA was isolated from the prepupal legs of 10 individuals using RNAqueous™ Total RNA Isolation Kit (Invitrogen). cDNA was synthesized using the template-switching method (see [25] for details). Library samples of cDNA were run using on an Illumina HiSeq 2000 platform at the Beijing Genomics Institute for 100 cycles with paired-end reads. All post-processed reads were assembled using Trinity (ver. R20131110) [26] to generate a transcript catalog. Using the sequences from the red flour beetle (*Tribolium castaneum*) as queries, BLASTX searches were performed to identify the orthologs of *Dll*, *hth* and *dac* in S*. femorata.* Reciprocal BLASTX were then performed to confirm the gene homology. The used sequences are provided in Supplementary Material.

Double stranded RNA synthesis of limb-patterning genes

Gene specific primer for *Dll*, *hth*, and *dac* with T7 adopter sequence (see Supplementary material) were used to amplify template DNA for for double-stranded RNA (dsRNA) synthesis. cDNA reverse-transcribed from total RNA described above was used as the template for PCR amplification of *Dll*, *hth*, and *dac*. For the control green fluorescent protein (*gfp*) control, the pGLO plasmid (BioRad, 1660003JEDU) was used as the template and T7-ligated primers (see Supplementary Material). The PCR product for *gfp* was purified and concentrated using Wizard® SV Gel and PCR Clean-Up System (Promega Corp., Wisconsin, USA) by following [27]. dsRNA was synthesized for each target gene following [28] using the Ambion MEGAscript® T7 Transcription Kit (Thermo Fisher Scientific, Waltham, MA, USA).

### Knockdown of limb-patterning genes

To investigate the effects of knockdown (KD) of the appendage genes on structural color formation, final instar larvae were removed from kudzu stems and injected with 1 μg of dsRNA targeting *hth*, *dac*, *Dll* or *gfp*. Each larva was placed individually in a plastic vials containing a piece of tissue paper and were maintained in an incubator at 25 ± 2 oC. Two weeks after eclosion, the structural coloration of the KD individuals was visually assessed to determine whether it differed from that of wild type individuals (Supplementary Fig. 1). From the KD experiment, we found that only the structural color of *Dll* KD individuals exhibited a structural coloration difference from that of the wild type individuals.

To further investigate structural color formation in *Dll* KD individuals, changes in structural coloration after eclosion were compared between *Dll* KD and *gfp* KD individuals. Larvae were removed from kudzu stem and were individually placed in plastic vials with tissue paper. Within 24 h of pupation, 1 μg of *Dll* dsRNA was injected into the thorax of each pupa (n = 14) using a 10 μL microsyringe (Hamilton Corp., Nevada, USA). As a control, 16 individuals were individually injected with 1μg of *gfp* dsRNA in the same manner.

### Transition of structural color formation

The structural color of *S. femorata* changes after eclosion and the color stabilizes by the sixth day after eclosion (Katsuki, personal observation). To investigate changes in structural color after eclosion, the wavelength of peak reflectance was measured for six days from the second to the seventh day after eclosion. The wavelength corresponding to the highest reflectance peak in the thorax was used as an indicator of the structural color. To measure the peak reflectance wavelength, each individual was placed on a 90 mm petri dish on the measurement stage. A probe (R200-7-UV/VIS, Ocean Insight, Inc., USA) was fixed vertically above the stage and connected to a spectrometer (FLAME-S-UV-VIS, Ocean Insight, Inc., USA) and a halogen light source (HL-2000-LL, Ocean Insight, Inc., USA) to measure reflectance across wavelengths. Reflectance from 350 to 800 nm was measured using OceanView software (Ocean Insight, Inc., USA). During the experiment, all individuals were maintained separately in plastic vials containing a piece of tissue paper.

### Observation of cuticular surface layers by SEM

Samples for scanning electron microscopy (SEM) observation were prepared as follows. One DAY 8 sample from each of *Dll* KD and *gfp* KD (control) treatments was stored in a freezer and subsequently dried in a desiccator over silica gel for two weeks. The elytra were removed from the body, and a part of elytra was excised for SEM observation. The samples were immersed in 2.5% glutaraldehyde in 0.1 M phosphate buffer and rinsed with 0.1 M phosphate buffer. The samples were then fixed with 1.5% osmium tetroxide in 0.1 M phosphate buffer and rinsed with distilled water. The fixed samples were dehydrated sequentially in ethanol solutions at the following concentrations: 30%, 50%, 60%, 70%, 80%, 95%, 100%, and 100% again. Each dehydration step lasted 10 - 30 min. The dehydrated samples were embedded in Spurr Low-Viscosity Embedding Media (Polysciences, Inc., Warrington, USA). The embedded samples were sectioned using an ultramicrotome (UltraCut UCT Type 706200, Leica, Munich, Germany) and coated with osmium using an osmium plasma coater (HPC-20, Vacuum Device Inc., Ibaraki, Japan). The cuticle region of the coated samples was observed using a scanning electron microscope (JSM-7900F, JEOL Ltd., Tokyo, Japan) at an accelerating voltage of 2 kV.

### Statistics

To assess whether the wavelength of peak reflectance differed between treatments, a robust linear mixed-effects model (robust LMM) was fitted using the rlmer function in the robustlmm package for R. Individual ID was included as a random effect while treatment, day, and their interaction were included as fixed effects. *P* values were calculated using Satterthwaite approximated degrees of freedom. Because a significant treatment × DAY interaction was detected in the robust LMM, simple-effects comparisons between treatments were conducted for each day using estimated marginal means (EMMs) derived from the robust LMM to examine treatment effects within each day. In addition, simple-effects tests of day within each treatment were conducted using EMMs to evaluate temporal changes within each treatment. Post hoc pairwise comparisons among treatment levels within each day and among day levels within each treatment were performed using the emmeans package with Tukey’s adjustment for multiple comparisons. All statistical analyses were performed using R version 4.5.2 [29].

## Results

### Transition of structural color formation

In control (*gfp* KD) individuals, the structural coloration of cuticle shifted from green to pink-purple, with wavelengths of peak reflectance of 580 ± 9 nm (mean ± SD) on DAY 2 (Fig. 1a), 667 ± 24 nm on DAY 4 (Fig. 1c) and 686 ± 7 nm on DAY 7 (Fig. 1f), respectively (Table 1; Fig. 1a-f). Within *gfp* KD individuals, significant differences in peak reflectance were detected among days (Tukey adjusted *P* < 0.05) expect between Day 3 and 4, Day 4 and 5, Day 5 and 6, and Day 6 and 7 (Table S1). In contrast, *Dll* KD individuals retained green structural coloration with a peak reflectance wavelength of 595 ± 11 nm (mean ± SD) on DAY2. The peak reflectance wavelength remained 591 nm to 596 nm throughout the post-eclosion development until DAY 7 (Table 1; Fig. 1g-l). No significant differences in peak reflectance wavelength were detected among days in *Dll* KD individuals (Tukey adjusted *P* > 0.05, Table S2). Peak reflectance wavelengths differed significantly between the two treatments throughout the observation period (Table S2; Fig. 2).

**Figure 1.**
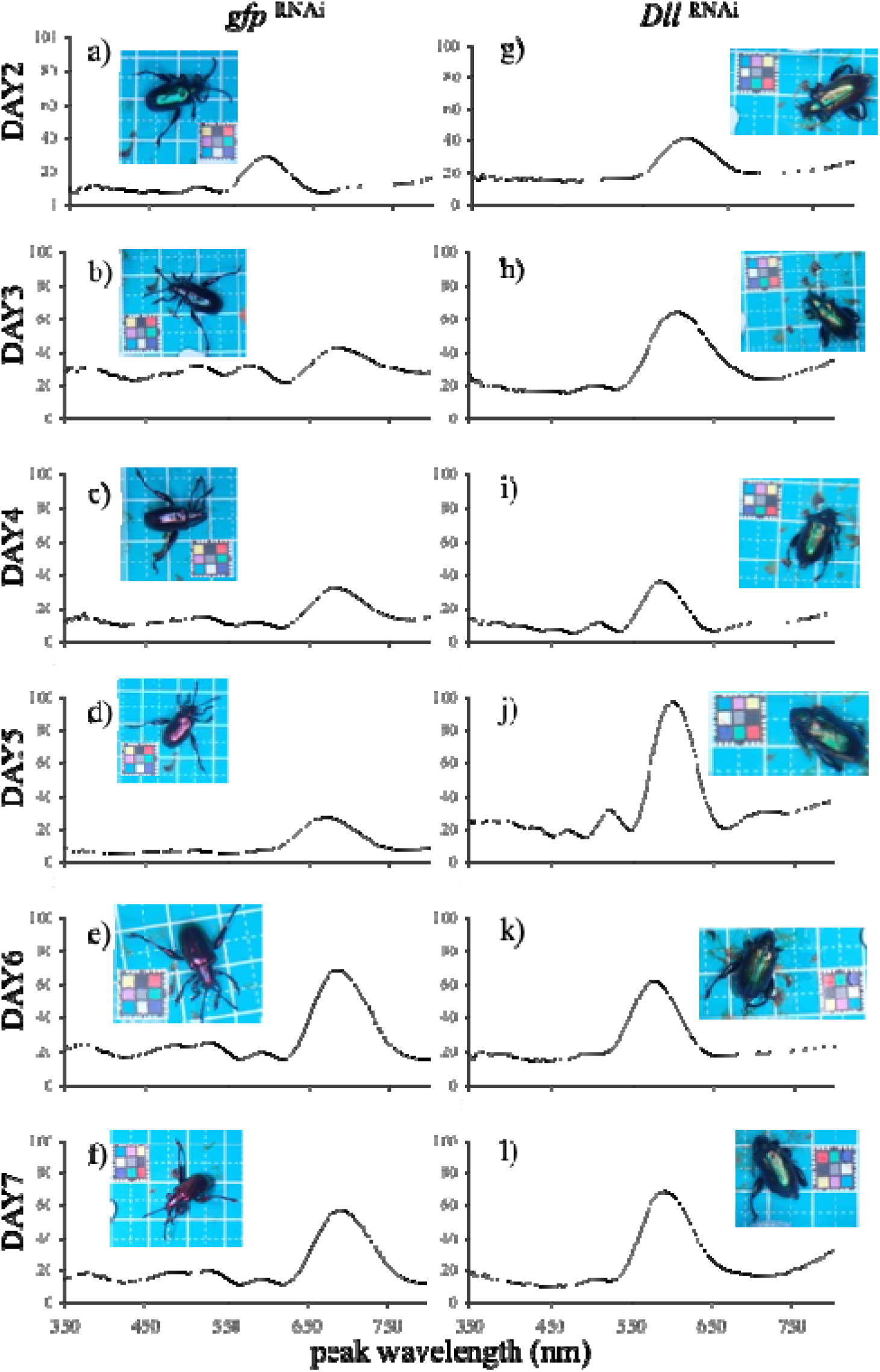
Transition of structural color (reflection wavelength, nm) from second day (DAY2) to seventh day (DAY7) after closing elytra in a-f) *gfp* ^RNAi^ (control) and g-l) *Dll* ^RNAi^ individuals. Note that some of the photographs have been edited to move the color chart closer to the sample, however, the chart used is taken from the same original image. Statistical analyses were performed using the full dataset. For clarity, only one individual per treatment is shown in this graph.

**Figure 2.**
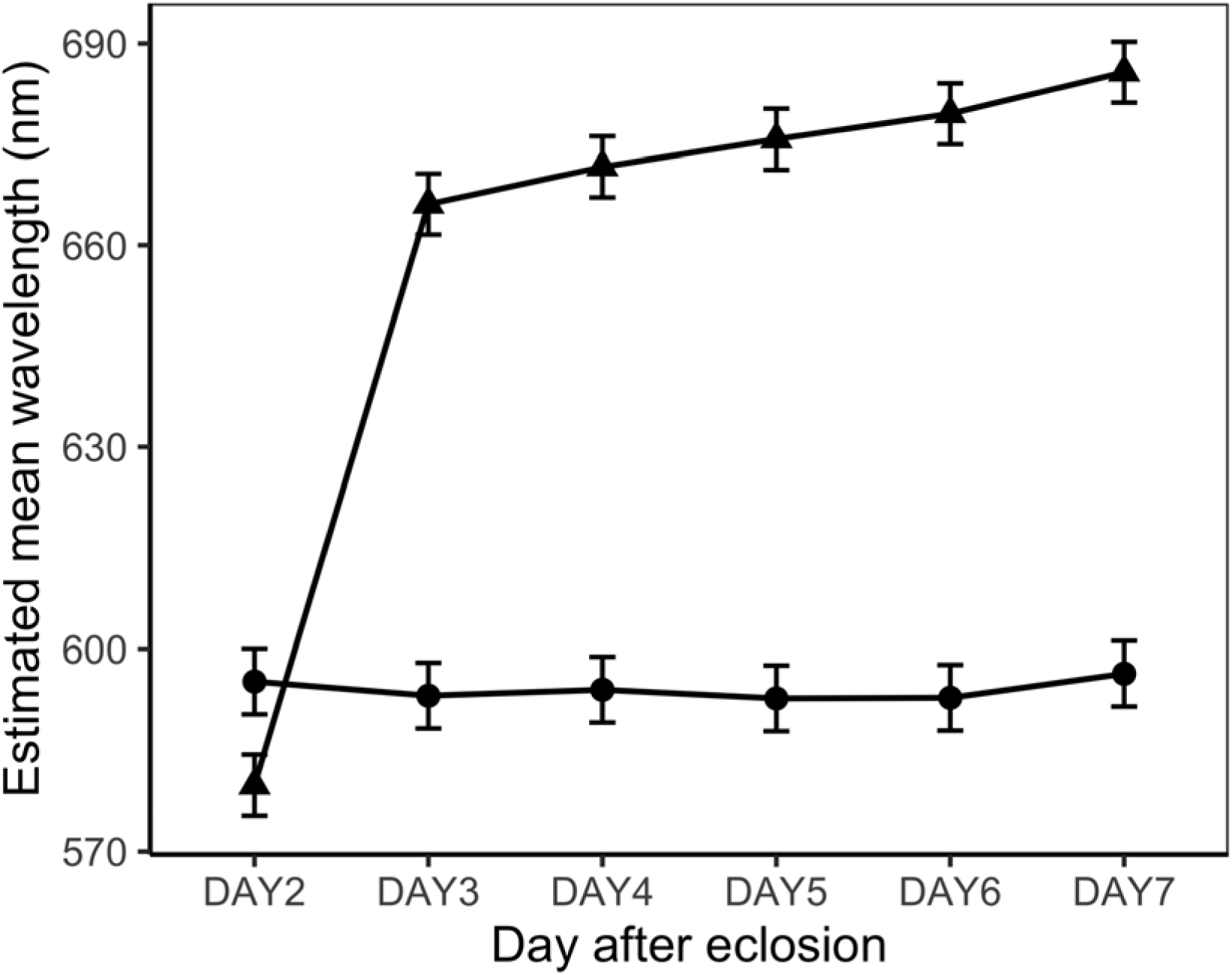
Estimated marginal means of peak of reflected wavelength (nm) obtained from a robust linear mixed model and the 95% asymptotic confidence intervals in each day. Triangle and circle points indicate *gfp* ^RNAi^ (control) and *Dll* ^RNAi^ treatments, respectively.

**Table 1.** Peak reflection wavelengths of *Dll* ^RNAi^ and control (*gfp* ^RNAi^) treatments.

| treatment | DAY | n | mean peak wavelength (nm) | s.d. |
|---|---|---|---|---|
| Dll | 2 | 14 | 595 | 11 |
|  | 3 | 14 | 592 | 10 |
|  | 4 | 14 | 594 | 9 |
|  | 5 | 14 | 592 | 10 |
|  | 6 | 14 | 592 | 14 |
|  | 7 | 14 | 596 | 9 |
| gfp | 2 | 16 | 580 | 9 |
|  | 3 | 16 | 639 | 51 |
|  | 4 | 16 | 667 | 24 |
|  | 5 | 16 | 676 | 4 |
|  | 6 | 16 | 680 | 6 |
|  | 7 | 16 | 686 | 7 |

### Cuticle layers in the elytra

The elytra surface of *S. femorata* exhibited hexagonal structures formed by surface irregularities, including spots and lines, and these surface structures did not appeared to be affected by *Dll* KD (Fig. 3a, d). In vertical sections, the procuticle was thinner in the *Dll* KD individuals than in control individuals (Fig. 3b, e). In the surface layers of the cuticle, two types of layers,dark and light, were laminated to form multilayer reflectors (Fig. 3c; f). These layers were located immediately beneath the cuticle surface and consisted of six or seven dark layers and six or seven light layers. The number of dark and light layers did not differ between treatments; however, the light layers were thinner and the dark layers were thicker in the *Dll* KD individual compared to the control.

**Figure 3.**
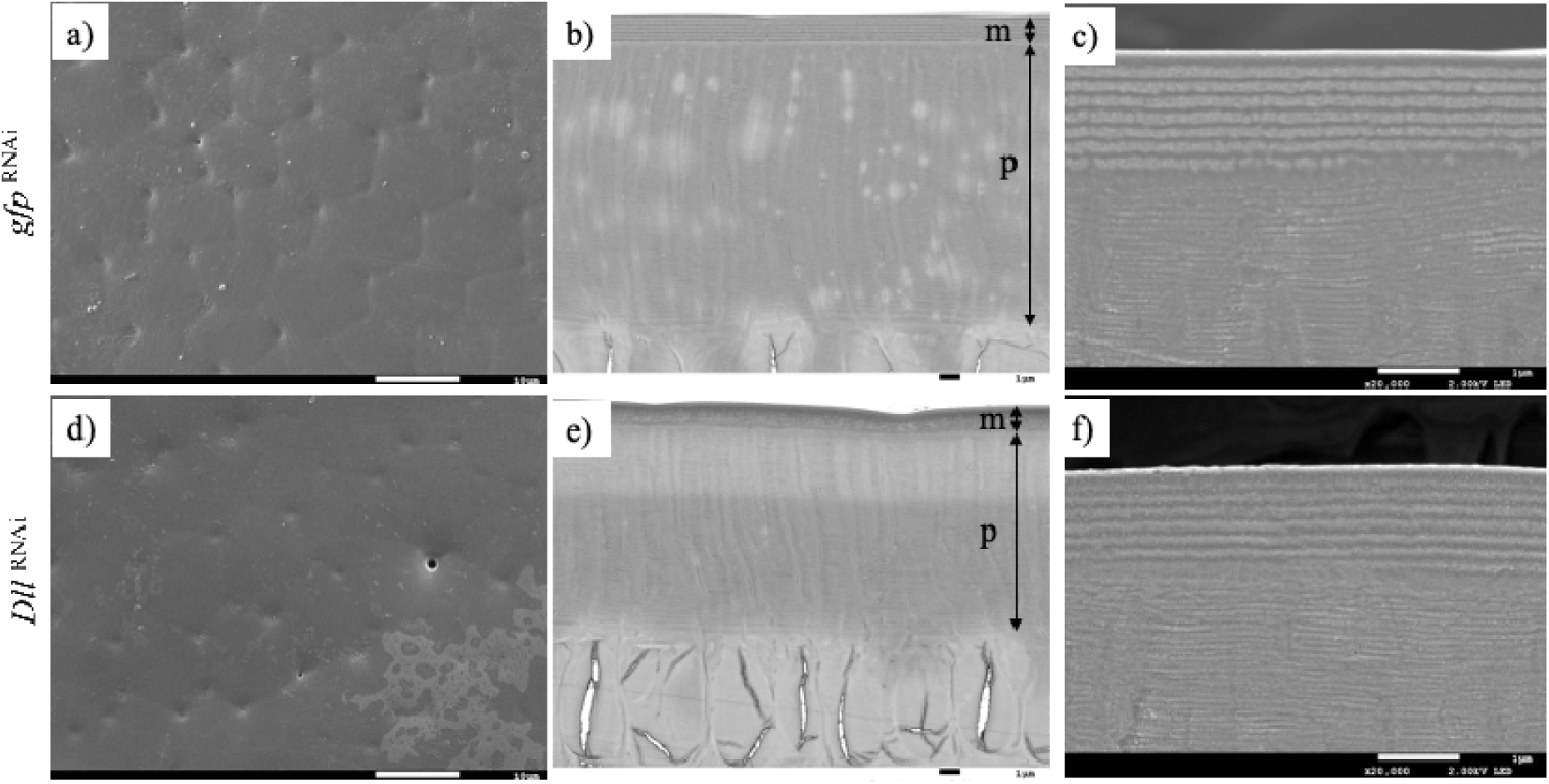
Microstructures of elytra in *gfp* ^RNAi^ treatment (a-c) and *Dll* ^RNAi^ treatment (d-f). Elytra surface (a and d), backscattered electron images of elytra dorsal cuticle (b and e) and secondary electron images of the reflectivelayer (c and f). External surfaces are uppermost (b, c, e and f). m) multiple reflector layers. p) procuticle. Scale bars are 10 μm (a and c), 1 μm (b, c, e and f), respectively.

## Discussion

We found that elytral cuticle of control *S. femorata* has multiple layers, consisting of thick light and thin dark layers, immediately beneath the elytra surface. This structure is consistent with previous reports of multiple interference structures in superficial regions of cuticle in other coleopteran species [20,21]. These findings suggest that these multilayers structures contribute to structural coloration in *S. femorata*. In insects, the cuticle develops and hardens through processes of cuticle melanization and sclerotization during pupation and immediately after eclosion [30, 31]. In *S. femorata*, elytral cuticle remains transparent during the pupal period and subsequently develops green structural coloration, which shifts to pink-purple after eclosion. During post-eclosion development, the wavelength of peak reflectance shifts from shorter wavelength (approximately 580 nm) to longer wavelength (approximately 685 nm), suggesting that the multilayer reflectors undergo structural changes during this period, potentially altering their optical properties and refractive index.

In contrast, *Dll* KD individuals didn’t exhibit a shift in structural coloration during post-eclosion development and retained green structural coloration with peak reflectance remaining at approximately 590 – 600 nm. The light layers were thinner and the dark layers were thicker in *Dll* KD cuticle than control cuticles, which may contribute to the difference in structural color. Indeed, in the jewel reed beetle *Plateumaris sericea,* individuals with thicker and with thinner reflector layers exhibit red and blue structural color, respectively, supporting the possible relationship between the thickness of the reflector units and the resulting structural color [21]. Aside from the multilayer reflectors immediately beneath the cuticle surface, the thickness of the procuticle beneath of these layers also differed between control and *Dll* KD cuticles. It is possible that these deeper layers also contribute to the difference in the structural coloration, for example, if pigments such as melanin within these layers absorb or reflect transmitted light [19]. Further observations will be needed as future studies to test this possibility.

Givin the limited information obtained from our SEM analysis, we refrain from concluding whether the multilayer reflectors in *S. femorata* are derived from the epicuticle or exocuticle. Many studies have reported multilayer interference structures in the epicuticle [15, 20, 21]. In general, insect cuticle is composed of chitin, proteins, lipids, and minerals whereas chitin is generally considered to be absent from the epicuticle [32, 33]. However, Lee et al. [15] has reported that the multilayer reflectors in the epicuticle of a jewel beetle contain chitin rich layers. Thus, multilayer reflectors in the may contain chitin, as do other cuticlar structures. Alternatively, it is also possible that multilayer reflectors belong to the exocuticle. Epicuticle completes its development at early stage of molting and eclosion but exocuticle continues. Onelli et al. [31] has concluded that the multilayer reflectors beneath the cuticle surface of the dock leaf beetle *Gastrophysa viridula* belong to the exocuticle based on observations of its developmental process. In *Tribolium castaneum*, which is a model Coleoptera species, the development of elytra cuticle has been well characterized. Epicuticle formation begins at 72h after the pupation and appears to be complete by 78h after pupation. On the other hands, procuticle deposition begins from approximately 78h after pupation and continues for several days after eclosion [34]. In our observations, the thickness of multilayer reflectors in wild type individuals at DAY 2 (green structural coloration; see Supplementary Fig. 2) differed from that at DAY 8, when structural coloration had developed fully. The light and dark layers in DAY 2 appeared more similar to those of the *Dll* KD individuals at DAY 7, which exhibited green structural coloration, than those of (control individuals at DAY 7, which exhibited pink-purple structural coloration. Collectively, these results suggest multilayer reflectors continue to develop during post-eclosion cuticle maturation, supporting the possibility of an exocuticular origin, as reported for *G. viridula*. Detailed observations of cuticul development throughout pupation and eclosion will be necessary to determine the precise origin of the multilayer reflectors in *S. femorata*.

*Distal-less* is a homeotic gene that function as a transcription factor for identifying the distal region of appendages, including legs and antennae [35]. When *Dll* was knocked down during late larval stage in *T. castaneum*, adult legs became shorter and the joints were fused [36, 37]. On the other hands, in a Lepidopteran species *Bicyclus anynana*, two eyespot patterns on the forewings are correlated with dynamic progression of the *Dll* expression. Overexpression of the *Dll* gene in larval stage induced increased eyespot size, whereas knockdown of *Dll* decreased it [38]. Moreover, ectopic expression of *Dll* induced ectopic patches of black scales. In contrast, depletion of *Dll* expression increased eyespot size and changed pigmentation in *Vanessa cardui* [39]. Consistently, *Dll* expression was correlated with the intensity of pigmentation in wing spot of *Drosophila biarmipes* [40]. These suggest that *Dll* functions broadly in epidermal patterning rather than solely in pigmentation. Because structural coloration is also generated through specialized differentiation and organization of the cuticle, *Dll* may contribute to structural color formation by regulating cuticle patterning and differentiation.

In Coleoptera, cuticle melanization, pigmentation and sclerotization, are largely associated with the tyrosine-mediated cuticle tanning pathway [33, 41]. In this pathway, tyrosine is converted to dopa- and dopamine-melanin and NADA- and NBAD-pigment, resulting in cuticle malanization and sclerotization. Beyond pigmentation, cuticle tanning also affects the physical properties of the cuticle. Structural coloration, however, depends primarily on the precise organization and optical properties of cuticular multilayers rather than on pigments themselves. Therefore, genes involved in cuticle differentiation may also influence structural coloration. Our results raise the possibility that *Dll* regulates aspects of cuticle differentiation that are important for formation of structural coloration. Because structural coloration depends on the precise organization and material properties of cuticular reflective layers, *Dll* may contribute to their development by regulating cuticle differentiation. Further studies, including transcriptome comparisons between *Dll* KD and control individuals, will be necessary to clarify the relationship between *Dll* and cuticle development, melanization, sclerotization and structural color formation in *S. femorata*.

## Data Accessibility Statement

The source data are provided in electronic supplementary material, Data S1.

## Declaration of AI use

AI tools (ChatGPT, OpenAI; GPT-5.6 Sol) was used to assist with troubleshooting analysis scripts, and with English language editing and improving the clarity of the text. All AI-assisted outputs were reviewed and revised by the authors, who take full responsibility for the final content of the manuscript.

## Competing Interests Statement

We declare that there are not any competing interests.

## CRediT authorship contribution statement

Masako Katsuki: Writing – review & editing, Writing – original draft, Formal analysis, Funding acquisition, Data curation, Investigation, Methodology, Conceptualization. Takashi Matsuo: Writing – review & editing. Conceptualization. Ken Ishikawa: Writing – review & editing, Formal analysis, Methodology. Yasukazu Okada: Writing – review & editing, Writing – original draft. Manabu Kamimura: Writing – review & editing, Conceptualization.

## Acknowledgement

We thank to JEOL Ltd. for technical support in SEM observation. This work was supported by Japan Society for the Promotion of Science (Grants-in-Aid for JSPS Fellow 19J40100 and 26KJ0157 and KAKENHI 22K06387) and Hokuto Foundation for Bioscience to MK.

## Notes

### Competing Interest Statement

The authors have declared no competing interest.

## References

1. Johansen, V.E., Catón, L., Hamidjaja, R., Oosterink, E., Wilts, B.D., Rasmussen, T.S., Sherlock, M.M., Ingham, C.J. and Vignolini, S., 2018. Genetic manipulation of structural color in bacterial colonies. Proc. Natl. Acad. Sci. 115(11), 2652–2657. 10.1073/pnas.1716214115

2. Kientz, B., Vukusic, P., Luke, S. and Rosenfeld, E., 2012. Iridescence of a marine bacterium and classification of prokaryotic structural colors. Appl. Environ. Microbiol.78(7), 2092–2099. 10.1128/AEM.07339-11

3. Kientz, B., Agogué, H., Lavergne, C., Marié, P. and Rosenfeld, E., 2013. Isolation and distribution of iridescent *Cellulophaga* and other iridescent marine bacteria from the Charente-Maritime coast, French Atlantic. Syst. Appl. Microbiol. 36(4), 244–251. 10.1016/j.syapm.2013.02.004

4. Vignolini, S., Moyroud, E., Glover, B.J. and Steiner, U., 2013. Analysing photonic structures in plants. J. R. Soc. Interface, 10(87), 20130394. 10.1098/rsif.2013.0394

5. Seago, A.E., Brady, P., Vigneron, J.P. and Schultz, T.D., 2009. Gold bugs and beyond: a review of iridescence and structural colour mechanisms in beetles (Coleoptera). J. R. Soc. Interface 6(suppl_2), S165–S184. 10.1098/rsif.2008.0354.focus

6. Mäthger, L.M., Denton, E.J., Marshall, N.J. and Hanlon, R.T., 2009. Mechanisms and behavioural functions of structural coloration in cephalopods. J. R. Soc. Interface, 6(suppl_2), S149–S163. 10.1098/rsif.2008.0366.focus

7. Osorio, D. and Ham, A.D., 2002. Spectral reflectance and directional properties of structural coloration in bird plumage. J. Exp. Biol. 205(14), 2017–2027. 10.1242/jeb.205.14.2017

8. Cai, C., Tihelka, E., Pan, Y., Yin, Z., Jiang, R., Xia, F. and Huang, D., 2020. Structural colours in diverse Mesozoic insects. Proc. R. Soc. B: Biol. Sci. 287, 1930. 10.1098/rspb.2020.0301

9. Ghiradella, H., 1994. Structure of butterfly scales: patterning in an insect cuticle. Microsc. Res. Tech. 27(5), 429–438. 10.1002/jemt.1070270509

10. Prum, R.O., Quinn, T. and Torres, R.H., 2006. Anatomically diverse butterfly scales all produce structural colours by coherent scattering. J. Exp. Biol. 209(4), 748–765. 10.1242/jeb.02051

11. Tanaka, G., Taniguchi, H., Maeda, H. and Nomura, S.I., 2010. Original structural color preserved in an ancient leaf beetle. Geol.38(2), 127–130. 10.1130/G25353.1

12. Kinoshita, S., Yoshioka, S. and Kawagoe, K., 2002. Mechanisms of structural colour in the Morpho butterfly: cooperation of regularity and irregularity in an iridescent scale. Proc. R. Soc. London. Series B: Biol. Sci. 269(1499), 1417-1421. 10.1098/rspb.2002.2019

13. Kinoshita, S., Yoshioka, S. and Miyazaki, J., 2008. Physics of structural colors. Rep. Prog. in Phys. 71(7), 076401. 10.1088/0034-4885/71/7/076401

14. Wang, D., Liu, Z., Wang, H., Li, M., Guo, L.J. and Zhang, C., 2023. Structural color generation: from layered thin films to optical metasurfaces. Nanophotonics, 12(6), 1019–1081. 10.1515/nanoph-2022-0063

15. Lee, J., Terlier, T., Jang, Y.J., Lee, K.B. and Lee, Y., 2020. Structural colors and physical properties of elytra in the jewel beetle, *Chrysochroa fulgidissima*, using surface analytical techniques. Surf. Interface Anal. 52(10), 656–664. 10.1002/sia.6807

16. Wilts, B.D., Michielsen, K., Kuipers, J., De Raedt, H. and Stavenga, D.G., 2012. Brilliant camouflage: photonic crystals in the diamond weevil, Entimus imperialis. Proc. R. Soc. B: Biol. Sci. 279(1738), 2524-2530. 10.1098/rspb.2011.2651

17. Anderson, T.F. and Richards Jr, A.G., 1942. An electron microscope study of some structural colors of insects. J. Appl. Phys., 13(12), 748–758. 10.1063/1.1714827

18. Vukusic, P., Hallam, B. and Noyes, J., 2007. Brilliant whiteness in ultrathin beetle scales. Science, 315(5810), 348–348. 10.1126/science.1134666

19. Parker, A.R., Mckenzie, D.R. and Large, M.C., 1998. Multilayer reflectors in animals using green and gold beetles as contrasting examples. J. Exp. Biol. 201(9), 1307–1313. 10.1242/jeb.201.9.1307

20. Akamine, M., Ishikawa, K., Maekawa, K., Kon, M.. 2011. The physical mechanism of cuticular color in *Phelotrupes auratus* (Coleoptera, Geotrupidae). Entomol. Sci. 14(3), 291–296. 10.1111/j.1479-8298.2011.00448.x

21. Kurachi, M., Takaku, Y., Komiya, Y. and Hariyama, T., 2002. The origin of extensive colour polymorphism in *Plateumaris sericea* (Chrysomelidae, Coleoptera). Naturwissenschaften, 89(7), 295–298.10.1007/s00114-002-0332-0

22. Noh MY, Muthukrishnan S, Kramer KJ, et al. 2017. Development and ultrastructure of the rigid dorsal and flexible ventral cuticles of the elytron of the red flour beetle, *Tribolium castaneum*. Insect biochemistry and molecular biology 91: 21–33. 10.1016/j.cois.2016.05.004

23. Charles, J. P. 2010. The regulation of expression of insect cuticle protein genes. Insect biochemistry and molecular biology, 40(3), 205–213. 10.1098/rspb.2020.0301

24. Zhu, K.Y., Merzendorfer, H., Zhang, W., Zhang, J. and Muthukrishnan, S., 2016. Biosynthesis, turnover, and functions of chitin in insects. Annu. Rev. Entomol. 61(1), 177–196. 10.1146/annurev-ento-010715-023933

25. Aird, S.D., Watanabe, Y., Villar-Briones, A., Roy, M.C., Tedara, K., Mikheyev, A.S., 2013. Quantitative high-throughput profiling of snake venom gland transcriptomes and proteomes (*Ovophis okinavensis* and *Protobothrops flavoviridis*). BMC Genom.14: 790. 10.1186/1471-2164-14-790

26. Grabherr, M.G., Haas, B.J., Yassour, M., Levin, J.Z., Thompson, D.A., Amit, I., Adiconis, X., Fan, L., Raychowdhury, R., Zeng, Q. and Chen, Z., 2011. Full-length transcriptome assembly from RNA-Seq data without a reference genome. Nat. Biotechnol. 29(7), 644–652. 10.1038/nbt.1883

27. Ozawa, T., Mizuhara, T., Arata, M., Shimada, M., Niimi, T., Okada, K., Okada, Y. and Ohta, K., 2016. Histone deacetylases control module-specific phenotypic plasticity in beetle weapons. Proc. Natl. Acad. Sci. 113(52), 15042–15047. 10.1073/pnas.1615688114

28. Sugiyama, M., Ozawa, T., Ohta, K., Okada, K., Niimi, T., Yamaguchi, K., Shigenobu, S. and Okada, Y., 2023. Transcriptomic and functional screening of weapon formation genes implies significance of cell adhesion molecules and female-biased genes in broad-horned flour beetle. PLoS genetics, 19(12), p.e1011069. 10.1371/journal.pgen.1011069

29. R Core Team (2024). R: A Language and Environment for Statistical Computing_. R Foundation for Statistical Computing, Vienna, Austria.

30. f S, Arakane Y, Noh MY, et al. (2022) Chitin in insect cuticle. In: Sugumaran, M. (Ed.) Insect Cuticle - Chitin, Catecholamine and Chemistry of Complexation. Advances in insect physiology. Elsevier, pp. 1–110.

31. Onelli, O.D., Kamp, T.V.D., Skepper, J.N., Powell, J., Rolo, T.D.S., Baumbach, T. and Vignolini, S., 2017. Development of structural colour in leaf beetles. Sci. Rep.7(1), 1373. 10.1038/s41598-017-01496-8

32. Sugumaran, M., 2022. Cuticular sclerotization in insects–A critical review. Adv. Insect Physiol. 62, 111–214. 10.1016/bs.aiip.2022.02.001

33. Muthukrishnan, S., Mun, S., Noh, M. Y., Geisbrecht, E. R. and Arakane, Y., 2020. Insect cuticular chitin contributes to form and function. Curr. Pharm. Des. 26(29), 3530–3545. 10.2174/1381612826666200523175409

34. Noh, M.Y., Muthukrishnan, S., Kramer, K.J. and Arakane, Y., 2016. Cuticle formation and pigmentation in beetles. Curr. Opin. Insect Sci. 17, 1–9. 10.1016/j.cois.2016.05.004

35. Chen, B., Piel, W.H. and Monteiro, A., 2016. Distal less homeobox genes of insects and spiders: genomic organization, function, regulation and evolution. Insect Sci. 23(3), 335–352. 10.1111/1744-7917.12327

36. Angelini, D.R., Smith, F.W. and Jockusch, E.L., 2012. Extent with modification: leg patterning in the beetle *Tribolium castaneum* and the evolution of serial homologs. G3: Genes| Genomes| Genetics, 2(2), 235-248. 10.1534/g3.111.001537

37. Suzuki, Y., Squires, D.C. and Riddiford, L.M., 2009. Larval leg integrity is maintained by Distal-less and is required for proper timing of metamorphosis in the flour beetle, Tribolium castaneum. Develop. Biol. 326(1), 60–67. 10.1016/j.ydbio.2008.10.022

38. Monteiro, A., Chen, B.I.N., Ramos, D.M., Oliver, J.C., Tong, X., Guo, M.I.N., Wang, W.K., Fazzino, L. and Kamal, F., 2013. Distal L ess Regulates Eyespot Patterns and Melanization in *Bicyclus Butterflies*. J. Exp. Zool. Part B: Mol. Dev. Evol. 320(5), 321–331. 10.1002/jez.b.22503

39. Zhang, L. and Reed, R.D., 2016. Genome editing in butterflies reveals that spalt promotes and Distal-less represses eyespot colour patterns. Nat. Commun. 7(1), 11769. 10.1038/ncomms11769

40. Arnoult, L., Su, K.F., Manoel, D., Minervino, C., Magriña, J., Gompel, N. and Prud’homme, B., 2013. Emergence and diversification of fly pigmentation through evolution of a gene regulatory module. Science, 339(6126), 1423–1426. 10.1126/science.1233749

41. Arakane, Y., Noh, M.Y., Asano, T. and Kramer, K.J., 2016. Tyrosine metabolism for insect cuticle pigmentation and sclerotization. In Extracellular composite matrices in arthropods, pp. 165–220. Cham: Springer International Publishing.

