## Supplementary material for "Marked change of cuticular structural color caused by knocking down of *Distal-less* gene in a leaf beetle"

Supplementary material 1

Gene sequences and the gene specific primers, which are *Dll*, *hth* and *dac*, used in RNAi experiment in *Sagra femorata*.

>Sag_Dll

ATGTCGGCGGAGGGACACATCGGTCCGCCGACGCCACACGAATCTAATAATTCCACGCCG

GTATCCAAGAGCGCTTTCATCGAGCTGCAGCAGCATGGATACGCCCCCCTTAGGACTACG

TATCAGCACCATTTTAATTCGCCGGCTGGGAACGCGCATTCTGGTCCTACGGGTGGACAC

GACACCGGCTTCCCTAGTCCTAGGGGCGCACTCGGAGCTTACCCTTTTCCGCCCATGCAA

CAGCACTCTTACACCGGATACCATCTCGGATCTTATGCCTCCAACTGCCCACCCTCCCCG

AAAGATGACGAGAAATGCCTTGCTCTGGAACGGCCCGCGGGTGGTGGCAAGGGCAAGAAA

ATGCGGAAACCTAGAACCATCTACTCCAGTTTGCAACTGCAACAGTTGAACAGGCGTTTC

CAGCGAACGCAGTACCTGGCATTACCCGAACGGGCCGAACTGGCGGCCAGTTTAGGTCTC

ACGCAAACACAGGTGAAAATATGGTTTCAGAATCGCAGGAGTAAATATAAGAAGATGATG

AAGGCGGCGCAAGTCCCCGGCGGCACGAACAATAACGGTACTCCGGGCGCTCACTCAGGT

CTATTAGGTGCTAGCGGGAATTTGCCCTCGAACAGTCCGGGACCGCAGGGCCAGAATATC

ATGCAAGGTAGGATCAAAATGGAAGGTGGTGGTTCATCTAGCGGGAGCCCTGTGGGAGGT

GCGGGAGCCGGCGGTTACCTGCAACATCCGTCACCCGCGCCCTCCTCCACTCCCGGTTCG

GATATGAGTGGCGGTGGCGGACATCCCGGAGCCTCGGGCAGTCCCGCCTCGGTGGTACAC

GGATGGGGCGACGTGAAACCACCTTCGGGAGCGGGGCCAGGACAACAGCAACAGCCGCCA

CCTACGCATCCCCATCCACACCACCCGCATCATCCGCCTCCCCCGCCCCCTACGCACAAT

TACCTGCCGCAGTACTCGTGGTATCCAGATAATCCGGGACTCCTCACGGTATGGCCAGCT

GTTTAA

>Sag_hth

ATGGAGTGCGGAGTTCAAAGCGCGATTCTGAGTGTACCTCCGGTGGGAGACGCCGCCGTG

CAACAGCCATTGCCCGCAGCTTATTCTCCGTCCGAATCGGAACAGGCACAATTCGAGGCC

GACAAAAGAGCTGTTTACAAGCATCCCCTATTCCCTCTGTTAGCGTTGCTTTTTGAAAGA

TGCGAATTGGCGACCCAATCGTCGGATACCCAGTCTAGTGACGCTTTTAATTTAGATATT

CAAGCTTTTGTGCAACACCAAGAAAGAGATAGGAAGCCTTTTTTAGCAAATGAACCAGAA

GTCGATGGACTGATGATAAAAGCAATTCAAGTGTTGAGGATACATCTTTTGGAATTAGAG

AAAGTTCAGGAACTATGTAAAGATTTCTGTAACCGATATATTACATGTTTAAAAGGAAAA

ATGCAATCAGAAAATTTGTTACGATCGGACTATCCACAAGATGTTTTAGGGACTGGTTTA

TTAAATAATATGGAAAATAATAATAACATGATGTCACATGGAAATTCAGAAAGTGGATCG

GCATCAAATAGTCCTGTACAATATTCGATAAGTCCATCACAACCGCCTGTGATTCCTAAT

CAAGCAAACCTGATAGACCGCCCTATAGAACAGAATAACTACCCTATTCCTCATTACGGG

GGTCCAACACCTCCTTTACCACCATCACAACCGCCCACAGAAAGTCTGGTGGTACATGGT

TCCACACCGTTATCACAAATAGGTAGCAATATGTGTTCAGTTCCTGCAAGCAGCGAAGGC

TCTGGGGCTCCTTTGTCGCCTGCGCCTACAACTCCTGGAGGCTCCGCCGATGATTGTGAC

AGTGATTTTAACTCCCGAGGGGGTCGGAAACAAAAACGAGGTGTTCTGCCTAAACATGCA

ACTAGTGTGATGCGGTCATGGCTTTTTCAACATTTAGTGCATCCTTATCCAACAGAAGAC

GAAAAACGCCATATAGCTGCCCAAACTAACCTAACACTTTTACAAGTGAATAATTGGTTT

ATCAATGCAAGACGAAGAATCTTGCAGCCAATGTTAGATGCTAGTGCACCAACTGATAAC

AATAGCACTACTAATTCGCCCAATCATAAGAAAAAGAAAACACAAAATAATAATAGGAGA

TTCTGGTCACAGGATTTCGCCAACATACAACCTCACTTGGATAGCAGTATTCTTTCAAGT

TCTGAAGATGAAGGGGATTCCTATAGCAGTGACGGAGAAGGCAGTTTAGTAATAGATGCG

AACGCTCAAAGTGGAGAAGTTAACCAAAATGGGCATCCTCCAACAGTCGATGCTTTAGGA

TATAATGACAATTCAACAGATATTACTAACAGCAAAAGACAGTGA

>Sag_dac

ATGATGGACAATTCCGGTATGGATTCAGGTAGTGATCACAGTAGTCACGCGTCTTCAGGT

TCACCCGCTGTGGCAGTTAAACCTTCTCCGTCGCCTCCGAATAATAATAACAACAATCAC

GGCCACAGTCAGGTACACAGCCCGAGACAAGCGAGTCCACTTGGCCCTCCGATGACCATG

GCTCATCCGCTTAGCCCACCTCCAGTGTCTTCAGCAAATTTGGCCAGAAGCATGGCATTG

GGACATTTGCCACCACCTGGTCTTGGACTTCTTAATTCCTTAGGAGTCTTACACAATCCT

CTGGATTTAATGGCACATCACGCTCCACCTCGTTCTTACAACAGTCCTCCGCCGATATCT

ACTTCAGATCCTACGGCGAACGAGTGCAAATTAGTTGATTATCGTGGACAAAAAGTGGCA

GCATTTATTATTGCCGGGGACACTATGTTGTGCCTTCCGCAAGCTTTCGAACTGTTCTTG

AAACATCTCGTTGGTGGACTGCACACAGTGTATACCAAGTTAAAGAGACTAGACATTGTT

CCTCTTGTTTGCAACGTGGAACAAGTGAGAATCTTGAGAGGACTCGGTGCCATCCAACCA

GGTGTCAATAGATGCAAACTGCTCTCCTGTAAAGACTTCGACACATTGTACAGAGATTGT

ACGACAGCCAGTTCCAGGCCCGGTCGACCACCGAAGAGAGCGCCTGTAGGACTGAGTCTC

GCTGCTTCTCACCTACAGCATCAACAGCTCAAAAAACAGAGAATGGACAACGGCGAGTAT

CCATACGAAAACGGACATATGGGTGATATGTCTCGACTAGAAAAATCTCCACTTCTAGCC

AACGGTTACAATCACCCGCCTCATCTCAGTCACATGCAATTCATGCAACTGCCCCATCCT

GCAGCAGCGCACTCGGCGCTCTTATCACCTGCTATGCCTCATAATTTGGCGCGACCCGAC

GGGTCAGTTATTAAAAATCAAGGAATGCCCACGATGGAAGCCCTCGCAAGGTCAGGCATC

TGGGAAAATTGCAGAGCAGCGTATGAAGACATCGTCAAACATCTTGAAAGGTTACGAGAA

GAAAGGTGCGACCCAGACAGACCCATTCCGCTGGACCAGAAACCCAGGGATCTTAGTCCA

AGAAATGGTTCCCCAACTGACCACAGTCCAGTTCTAAATCTTTCAAAAAGTGGTGCAGGC

AGTGCTGGTTCAACAGGAGACGCAGATCATTCTGGTAGCGAGGCTGATGGTCCTGATGCT

CCTCCGACGCCTAGATCACCTCGATCGGCGGCTGAAGACGACGAGGATGACAATATCTCC

GAGGGAGAGGACGACGACGAAAAGGATCAAGATATGGACGACGGCGATCTTCCGCTTCCT

TCCGCCCCCAGCCACGATTCCCCTCATCATCACCAAGCCCTGAACTATTCTACTCTGGCT

AGCGCCATGGCCAGCGCCAACGGAGGAGCGGCGGGAGTGACCCCCACGGCTCCGCCGATG

GATCATTCGGGCGTTTCCTCAACCGAAACCCTACTCAGGAACATCCAAGGGCTGCTGAAA

GTGGCCGCTGATAACGCAAGACAGCAGGAGCGGCAAGTCAACTATGAAAAAGCTGAGCTC

AAGATGGATGTCCTAAGAGAACGAGAAGAGAAGGATAATTTAGAGAGACAATTGCTGGAA

GAACAAAAAATCAGAGTCATGGTTCAAAAAAGACTTCGAAGAGAAAAAAGACTCCGCAGA

AGAATACAACATCAATTAGAGATCGAAGCCAAACGACGTGCACAACTTGAAGAAGCTTTA

AGAGCCTCAGGTGCATCGGAACAAATTAGTATAATAAACGAAAAAATCGCTCAAGAATAT

CCTCTTCCTCCAACGACATCAATGAATGCCTCAACACATCCCCCATCTGCTTCTAGTGTG

TCCTCGAATCAACAACAAATCAATAATCCACCACCACCTCCTCCGGAACTGCGTCAACCT

GTAGGTATTGACAGGATTAAACAAGAACGCGAAGATACTTCTAGCACCCCTCAGGTGTAT

GGTGGTAGAGAGGCCCCACCGCCATCGGTACCGCCACCACAAGACACCAAACCCTGGGGC

TATGCGGGTATGGAAATGATGAGTACCGGGGCGGCCTTTTGGCAAAATTATTCCGATTCT

CTCGCACAGGAGCTGGAAATGGAGAGAAAATCAAGGCAGCAACAGGCCGAACGAGAAGTT

AAATCCCCATTACAAGACAGAGCAGCATATTACAAGAATTCTGTTCTGTTTACTAGCAGT

GCGACATAA

>Sag_Dll_forward

taatacgactcactatagggagCAAGAGCGCTTTCATCGAGC

>Sag_Dll_reverse

taatacgactcactatagggagATGATATTCTGGCCCTGCGG

>Sag_hth_forward

taatacgactcactatagggagCCATTGCCCGCAGCTTATTC

>Sag_hth_reverse

taatacgactcactatagggagGGCGGTCTATCAGGTTTGCT

>Sag_dac_forward

taatacgactcactatagggagGAAGAAAGGTGCGACCCAGA

>Sag_dac_reverse

taatacgactcactatagggagGCTGTCTTGCGTTATCAGCG

>gfp_forward

taatacgactcactatagggagCCTGTTCCATGGCCAACA

>gfp_ reverse

taatacgactcactatagggagAGGACAGGGCCATCGC
