## Supplementary figures and images for "Marked change of cuticular structural color caused by knocking down of *Distal-less* gene in a leaf beetle"

### Figure S1

Figure S1

Structural colors of a) *gfp* KD, b) *dac* KD, c) *hth* KD and d) *Dll* KD individuals.

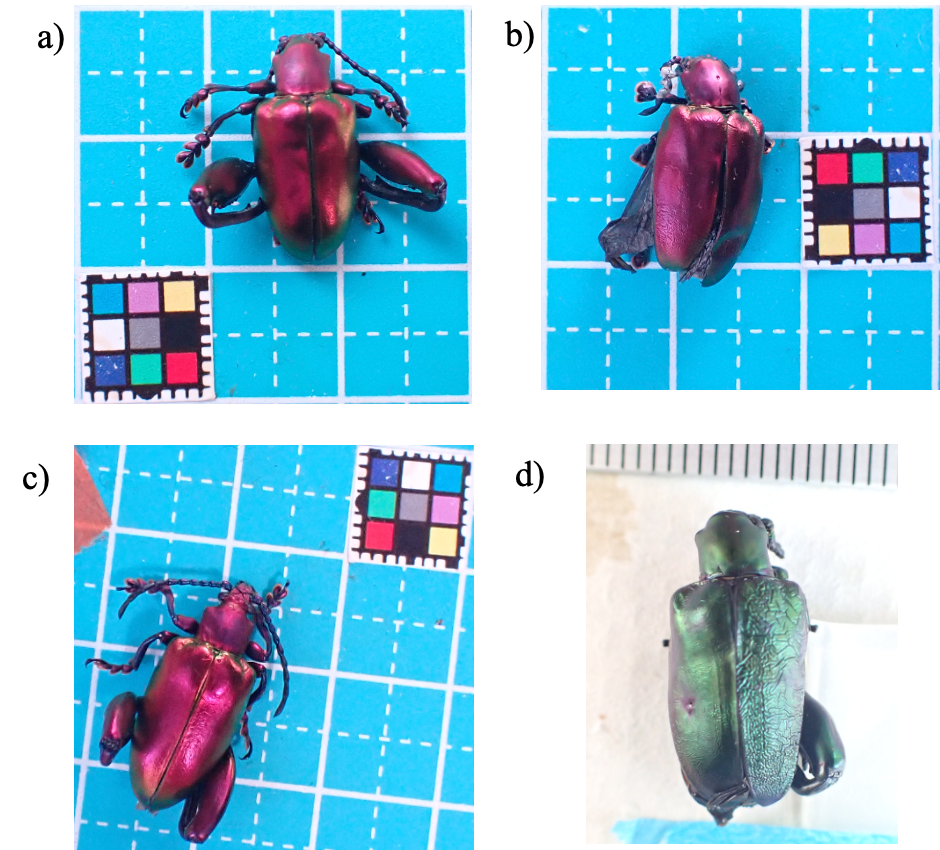
