## Supplementary material for "Marked change of cuticular structural color caused by knocking down of *Distal-less* gene in a leaf beetle": Figure S2

Figure S1

Microstructures of elytra ay DAY. 2 (next day from the eclosion) in wild type (non dsRNA injection) individual. Secondary electron images of the elytra cuticle including reflective layers.

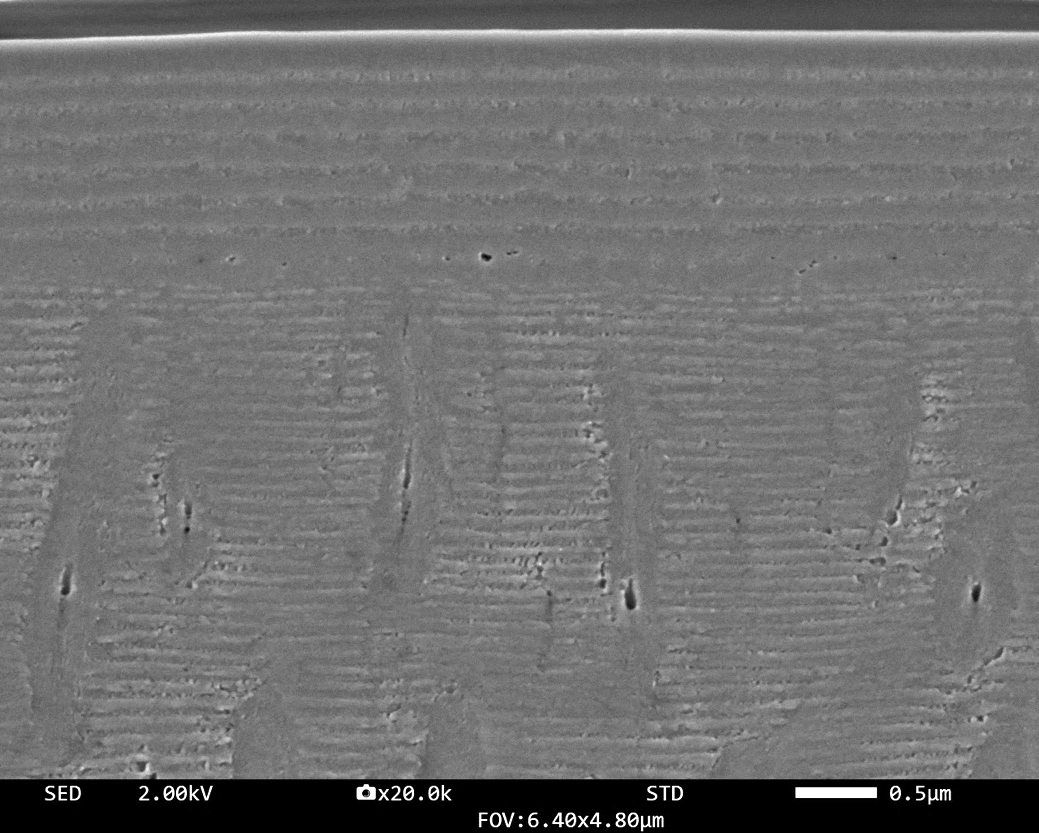
