## Supplementary material for "Marked change of cuticular structural color caused by knocking down of *Distal-less* gene in a leaf beetle": Table S1S2

| Table S1. Results of pairwise comparisons between days within a treatment with Tukey’s adjustment | | | | |
| --- | --- | --- | --- | --- |
| treatment | contrast | estimate | 95% CI | p-value |
| gfp | DAY2 - DAY3 | -86.266 | -93.987 − -78.544 | **<0.001** |
|  | DAY2 - DAY4 | -91.804 | -99.526 − -84.083 | **<0.001** |
|  | DAY2 - DAY5 | -95.904 | -103.626 − -88.183 | **<0.001** |
|  | DAY2 - DAY6 | -99.676 | -107.398 − -91.955 | **<0.001** |
|  | DAY2 - DAY7 | -105.870 | -113.592 − -98.149 | **<0.001** |
|  | DAY3 - DAY4 | -5.539 | -13.260 − 2.183 | 0.317 |
|  | DAY3 - DAY5 | -9.639 | -17.360 − -1.917 | **0.005** |
|  | DAY3 - DAY6 | -13.411 | -21.132 − -5.689 | **<0.001** |
|  | DAY3 - DAY7 | -19.605 | -27.326 − -11.883 | **<0.001** |
|  | DAY4 - DAY5 | -4.100 | -11.822 − 3.621 | 0.656 |
|  | DAY4 - DAY6 | -7.872 | -15.594 − -0.151 | **0.043** |
|  | DAY4 - DAY7 | -14.066 | -21.788 − -6.345 | **<0.001** |
|  | DAY5 - DAY6 | -3.772 | -11.493 − 3.949 | 0.732 |
|  | DAY5 - DAY7 | -9.966 | -17.688 − -2.245 | **0.003** |
|  | DAY6 - DAY7 | -6.194 | -13.916 − 1.527 | 0.200 |
| Dll | DAY2 - DAY3 | 2.089 | -6.165 − 10.344 | 0.979 |
|  | DAY2 - DAY4 | 1.211 | -7.044 − 9.466 | 0.998 |
|  | DAY2 - DAY5 | 2.496 | -5.759 − 10.750 | 0.956 |
|  | DAY2 - DAY6 | 2.401 | -5.853 − 10.656 | 0.962 |
|  | DAY2 - DAY7 | -1.153 | -9.408 − 7.101 | 0.999 |
|  | DAY3 - DAY4 | -0.878 | -9.133 − 7.376 | 1.000 |
|  | DAY3 - DAY5 | 0.407 | -7.848 − 8.661 | 1.000 |
|  | DAY3 - DAY6 | 0.312 | -7.943 − 8.567 | 1.000 |
|  | DAY3 - DAY7 | -3.242 | -11.497 − 5.012 | 0.874 |
|  | DAY4 - DAY5 | 1.285 | -6.970 − 9.539 | 0.998 |
|  | DAY4 - DAY6 | 1.190 | -7.064 − 9.445 | 0.999 |
|  | DAY4 - DAY7 | -2.364 | -10.619 − 5.890 | 0.965 |
|  | DAY5 - DAY6 | -0.095 | -8.349 − 8.160 | 1.000 |
|  | DAY5 - DAY7 | -3.649 | -11.903 − 4.606 | 0.807 |
|  | DAY6 - DAY7 | -3.554 | -11.809 − 4.700 | 0.824 |

| Table S2. Results of pairwise comparisons between treatments with Tukey’s adjustment | | | | |
| --- | --- | --- | --- | --- |
| treatment | contrast | estimate | 95% CI | p-value |
| Dll_gfp | DAY2 | 15.345 | 8.701 − 21.991 | **<0.001** |
|  | DAY3 | -73.010 | -79.651 − -66.371 | **<0.001** |
|  | DAY4 | -77.670 | -84.311 − -71.031 | **<0.001** |
|  | DAY5 | -83.055 | -89.701 − -76.411 | **<0.001** |
|  | DAY6 | -86.732 | -93.371 − -80.091 | **<0.001** |
|  | DAY7 | -89.372 | -96.011 − -82.731 | **<0.001** |
